# The dual orexin/hypocretin receptor antagonist suvorexant reduces addiction-like behaviors for the opioid fentanyl

**DOI:** 10.1101/2020.04.25.061887

**Authors:** Shayna L. O’Connor, Jennifer E. Fragale, Morgan H James, Gary Aston-Jones

**Author notes:** Corresponding Authors: Gary Aston-Jones, Ph.D., 683 Hoes Lane West, Piscataway NJ 08854, E, Morgan H. James, Ph.D., 683 Hoes Lane West, Piscataway NJ 08854, E. Authors contributed equally (Shayna L. O’Connor, Jennifer E. Fragale).

## Abstract

The orexin (hypocretin) system is critical for motivated seeking of all drugs of abuse, including opioids. In 2019, the National Institute on Drug Addiction (NIDA) identified the orexin system as a high priority target mechanism for novel pharmacological therapies to treat opioid use disorder (OUD). Suvorexant (Belsomra™) is a dual orexin receptor 1/orexin receptor 2 (OxR1/OxR2) antagonist that is FDA-approved for the treatment of insomnia, and thus has the potential to be readily repurposed for the treatment of OUD. However, studies have yet to test the therapeutic potential of suvorexant with respect to reducing opioid-related behaviors. Accordingly, here we investigated the efficacy of suvorexant in reducing several addiction-relevant behaviors in fentanyl self-administrating rats. In rats with limited drug experience, suvorexant decreased motivation for fentanyl on a behavioral economics (BE) task. This effect was greatest in rats with the highest motivation for fentanyl. Suvorexant was even more effective at decreasing motivation for fentanyl following induction of a more robust addiction phenotype by intermittent access (IntA) self-administration of the opioid. Suvorexant also attenuated punished responding for fentanyl and reduced cued reinstatement in IntA rats. Suvorexant did not affect general locomotor activity or natural reward seeking, indicating that at the doses used here, suvorexant can be used to reduce drug seeking with limited sedative or off-target effects. Together, these results highlight the therapeutic potential of suvorexant, particularly in individuals with the severe OUD.

## Introduction

Over the last two decades, opioid use and abuse in the United States has risen at alarming rates, culminating in the declaration of a public health emergency. Much of this increase has been driven by use of synthetic opioids, including fentanyl, which now represents the primary contributor to opioid-related overdose deaths [1]. Opioid substitution therapy (OST) is the standard treatment for opioid use disorder (OUD), with methadone, buprenorphine and naltrexone being most common [2,3]. Despite being effective in preventing relapse and overdose, OST is broadly underutilized in part due to concerns with stigma associated with iatrogenic opioid dependence[3]. Thus, there is significant interest in the development of non-opioid alternatives to OST for the treatment of OUD.

To this end, the National Institute on Drug Abuse (NIDA) recently identified the orexin (hypocretin) system as among the highest priority target mechanisms for novel therapeutics for treating OUD [4]. Orexins A and B are excitatory neuropeptides that are produced exclusively by neurons in the lateral and perifornical hypothalamus and are important for a range of physiological processes, including opioid-related behaviors [5-8]. Orexin-expressing neurons are activated during opioid withdrawal [9], and their activity is predictive of opioid seeking behavior [10]. Moreover, higher numbers of orexin-expressing neurons are observed in the brains of human heroin users [11], as well as in rats that exhibit a strong addiction-like phenotype for fentanyl. [12] Orexin peptides bind to two G-protein coupled receptors (orexin receptor 1, OxR1; and orexin receptor 2, OxR2) that are widely distributed throughout the brain, including in key reward and arousal centers [13]. Evidence generally supports a functional dichotomy for OxR1 vs OxR2 signaling, whereby OxR1 signaling preferentially mediates reward behaviors and OxR2 signaling modulates arousal and sleep [5]. Consistent with this, systemic treatment with selective OxR1 antagonists, including SB334867 (SB), reduce economic demand and cued reinstatement for all drugs of abuse tested, including the opioids fentanyl and remifentanil [14,15], but have limited effects on arousal and sleep [16-18]. In contrast, selective OxR2 antagonists promote sleep [18], and were reported to have limited effect on cocaine seeking behavior [19]. Despite this apparent dichotomy, emerging evidence indicates that OxR2 signaling is important for some addiction behaviors, including opioid seeking, under certain conditions. For example, systemic administration of the OxR2 antagonist NBI-80713 reduces heroin self-administration in rats with a history of extended (12h/d) heroin self-administration [20], indicating that signaling at OxR2 might become important under conditions of heightened motivation for opioid. Taken together, these data indicate that compounds that block signaling at both OxR1 and OxR2 (dual orexin receptor antagonists; DORAs) might have stronger anti-addiction properties compared to single orexin receptor antagonists (SORAs), particularly in highly addicted individuals [21,22].

In 2014, the DORA (Belsomra™) was FDA-approved for treatment for insomnia [23]. When administered immediately prior to bedtime, suvorexant decreases latency to sleep onset and increases time spent sleeping [23], presumably due to its actions at OxR2. However, its actions at *both* OxR1 and OxR2 raise the possibility that it could be repurposed for use in populations with substance use disorders, including OUD [21,22]. To date, only a small number of studies has tested the efficacy of suvorexant on drug-seeking behaviors, with each focusing on psychostimulants. Two studies in rats indicated that suvorexant reduced the motivational properties of cocaine and improved cocaine-induced deficits in impulsive behavior [24,25], and a preliminary randomized clinical study indicated that suvorexant tended to reduce several relapse-related indices in patients with cocaine use disorder [26]. To date, there has been no preclinical or clinical reports on the efficacy of suvorexant to reduce OUD behaviors [21]. Thus, here we sought to examine the therapeutic effects of suvorexant in fentanyl self-administering rats. We report that suvorexant reduced increased demand elasticity (decreased motivation) for fentanyl without affecting low-effort fentanyl intake; these effects were most pronounced in rats with the strongest motivation for fentanyl. Moreover, following induction of a strong addiction phenotype by intermittent access (IntA) self-administration of fentanyl [12], suvorexant was more effective at increasing demand elasticity, and also attenuated compulsive responding for fentanyl and reduced cued reinstatement of extinguished fentanyl seeking. Taken together, these data indicate that suvorexant is effective at reducing the OUD behavioral phenotype, particularly in individuals with the severe cases of OUD.

## Materials and Methods

### Animals

Adult male Long Evans rats (250-300g) were received from Charles River Laboratories (Raleigh, NC). All rats were pair housed on a reverse 12:12 hour light/dark cycle in a temperature and humidity-controlled animal facility. Animals had ad libitum access to food and water throughout the experiments. All procedures were approved by Rutgers University New Brunswick Institutional Animal Care and Use Committee and conducted in accordance with the NIH Guide for the Care and Use of Laboratory Animals.

### Drugs

Fentanyl HCl powder (National Institute of Drug Abuse [NIDA] Drug Supply Program) was dissolved in 0.9% sterile saline to a concentration of 8 μg/ml. Suvorexant (NIDA Drug Supply Program) was dissolved in dimethyl sulfoxide (DMSO; 100%) through vortexing; 0, 10 or 30 mg/kg was injected at a fixed volume of 100 μl (ip) 30min prior to testing, as described in previous publications [27].

### Surgery

Two weeks after arrival, animals were anesthetized with an ip injection of ketamine (56.5 mg/kg) and xylazine (11.3 mg/kg), administered the analgesic, rimadyl (5 mg/kg, sc), and underwent intravenous catheter surgery as described previously [19]. During recovery, catheters were flushed daily with the antibiotic cefazolin (0.1 ml; 100 mg/ml) and heparin (0.1 ml; 100 U/ml) and this was continued throughout the experiment. Animals were allowed 1wk to recover prior to self-administration training.

### Self-Administration Training

Animals were trained to self-administer fentanyl on a 2h fixed ratio 1 (FR1) schedule in sound-attenuated operant chambers (Med Associates, St. Albans, VT, USA), as described previously. Drug availability was signaled by illumination of the house light, and an infusion of fentanyl (3.6 s infusion of 0.5 μg of fentanyl, iv) was elicited when the active lever was pressed. Drug infusions were paired with illumination of a white stimulus light above the active lever, and tone (78 dB, 2900 Hz), both of which were presented for the duration of the infusion. Inactive lever responses were recorded but did not result in infusions or cues. Each infusion was followed by a 20 s time out, which was signaled by the house light offset. The operant chambers were controlled by Med-PC IV software. Animals were trained for a minimum of 10 sessions (1 session per day) and to a criterion of >25 infusion for 3 consecutive days.

### Within-Session Threshold Procedure

Following self-administration training, animals’ economic demand for fentanyl was assessed on the behavioral economics threshold procedure (BE) recently described by our laboratory [16]. Briefly, in this 110min paradigm, animals responded on an FR1 schedule for fentanyl infusions except doses of fentanyl were decreased every 10min on a quarter logarithmic scale by adjusting the duration of the infusion (1.12, 0.63, 0.36, 0.20, 0.11, 0.06, 0.04, 0.02, 0.01, 0.006, 0.004 μg fentanyl per infusion). As in FR1 training, the house light signaled drug availability, and each drug infusion was accompanied by light and tone cues. During each infusion, the house light was extinguished, and any presses on either lever are recorded but had no effect. Notably, there were no timeout periods following each infusion. Animals were trained for a minimum of 6 sessions (1 session per day) and until demand parameters (Q_o_ and α values) differed by less than 25% across three consecutive sessions.

Demand curves were generated using a focused fitting approach, as previously described [16,28,29]. All data points up until two bins past the point of maximum price paid per mg of fentanyl (Pmax) were used to generate curve fits. This was done to account for unstable brain fentanyl concentrations due to the rapid elimination of fentanyl from the brain [30]. Using this method, two parameters, Q_o,_ and α, were estimated from demand curves; Q_o_ represents the theoretical level of preferred drug consumption when no effort is required, and α reflects the slope of the curve and is taken to be an inverse measure of motivation.

#### Experiment 1: Efficacy of systemic suvorexant administration on fentanyl demand in rats with limited drug access

We first determined the effect of suvorexant on fentanyl demand in rats with a limited history of fentanyl self-administration. Rats were trained to self-administer fentanyl on an FR1 schedule until meeting training criteria (for an average of 11.3 ±1.0d), and then tested for demand on the BE procedure, as described above. After achieving stability on the BE procedure (10.5 ±1.0d), rats were tested in subsequent sessions 30min following systemic treatment of suvorexant (0, 10 mg/kg or 30 mg/kg; ip). Testing was carried out in a within-subjects manner, and the order of treatment was counterbalanced. Test days were separated by at least two regular sessions, and continued until Q_o_ and α values were stable, i.e., within ± 25% of pre-treatment values (baseline); this allowed us to test and account for potential persistent effects of suvorexant on subsequent sessions [15,31].

#### Experiment 2: Efficacy of systemic suvorexant on fentanyl demand, punished responding, and cue-induced reinstatement of fentanyl seeking following intermittent access to fentanyl

Next, we tested the efficacy of suvorexant on several addiction-relevant indices in rats with a history of intermittent access (IntA) to fentanyl. IntA is a self-administration procedure that models the transition to a multiphenotypic addiction-like state in rats [12]. To confirm the induction of an addiction-like state in IntA rats, we compared their demand values to a short access (ShA) control group that was tested concurrently; we previously demonstrated that ShA does not result in an addiction-like state [12]. Rats were pseudorandomly assigned to IntA or ShA groups such that baseline α values did not differ between access groups.

### Intermittent and short access self-administration procedures

A subgroup of animals (n=8) from Experiment 1 underwent IntA training for fentanyl for 14d. IntA sessions were 6h in duration, during which 5min drug access periods (signaled by illumination of the houselight and extension of the levers) were separated by 25min no drug access time outs (houselight off, levers retracted). During drug access periods, animals could press on a FR1 schedule for a 3.6s infusion of fentanyl (0.5μg) that was paired with a light and tone cue; infusions were not followed by a timeout period [12]. ShA rats (n=8) were given continuous access to fentanyl on an FR1 schedule in 1h sessions. Active lever responses resulted in a 3.6s infusion of 0.5μg of fentanyl paired with light and tone were proceeded by a 20s time-out period signaled by termination of the house light. Both IntA and ShA groups were trained for 14 consecutive sessions.

### Effect of suvorexant on fentanyl demand following IntA

Following IntA training, rats were re-stabilized on the BE procedure, as above. Rats were then tested for demand following treatment with suvorexant (0, 10, 30mg/kg; ip) in a counterbalanced manner, as above. Suvorexant-test days were separated by at least 2 BE sessions.

### Effect of suvorexant on punished responding for fentanyl

Following demand testing, a subgroup of animals previously given IntA were tested for compulsive (footshock-punished) responding for fentanyl; some IntA animals lost catheter patency and instead were tested for extinction and reinstatement, described below. In the punished responding procedure, the amount of fentanyl infused was high during the first bin (1.12μg), enabling rats to quickly achieve their preferred brain fentanyl levels. The dose was decreased beginning in the second bin to 0.112μg per injection, and was held constant for the remainder of the session. Rats were trained on this procedure for a minimum of 3d and until responding was stable (<25% variability in responding over 3d; typically 4.8 ±1.0d). The following day, beginning in the third 10min-bin, footshock (0.5s) was delivered during each fentanyl infusion, and these increased in amplitude every 10min on a tenth-log_10_ scale: 0.13; 0.16; 0.20; 0.25; 0.32; 0.40; 0.50; 0.63; 0.79 milliamps. Shock resistance was characterized as the maximum cumulative charge in millicoulombs (mC) an animal self-administered in any one bin. Rats were tested for shock resistance (a measure of compulsive responding) 30min following treatment with suvorexant (0, 30mg/kg; ip) in a counterbalanced manner; each test day was separated by a minimum of 3d on a ‘no shock’ paradigm and until baseline stability was achieved.

### Extinction and cued reinstatement testing

Lever pressing was extinguished in daily 2h extinction sessions, whereby lever presses did not elicit drug infusions or cues. Animals remained on this procedure for a minimum of 7 sessions, and until the last 3 sessions had ≤ 25 active lever presses. During reinstatement tests, active lever responses were accompanied by light and tone cues, but no drug infusion. Rats were tested for cued reinstatement 30min following treatment with suvorexant (0, 10, 30mg/kg; ip) in a counterbalanced manner; test days were separated by a minimum of 2 extinction sessions where responding during these sessions met extinction criteria (≤ 25 active lever presses) [32,33]. Following cued reinstatement tests, rats were tested with suvorexant (0 and 30mg/kg; ip) during 2h extinction sessions. Rats were administered suvorexant 30min prior to testing in a counterbalanced manner. Test days were separated by 48h with no extinction sessions in between testing.

#### Experiment 3: Effect of suvorexant on general locomotion and natural reward seeking

Because of the known hypnotic properties of suvorexant when administered prior to the onset of the inactive period, we tested whether the doses used here were associated with impairments on i) a task requiring low-effort responding for a natural reward (sucrose), and ii) general locomotor reactivity to a novel environment. These experiments included rats from both *Experiment 1* and *Experiment 2*.

### Locomotor activity

Rats (n=19) were placed in novel locomotor chambers (clear acrylic, 42cm x 42cm x 30cm) equipped with SuperFlex monitors (Omintech Electronics Inc, Columbus, OH) containing a 16 × 16 infrared light beam array for the x/y axis (horizontal activity) and 16 × 16 infrared light beams for the z axis (vertical activity). Activity was recorded by Fusion SuperFlex software. Rats were pseudorandomly assigned to treatment groups (0 or 30mg/kg suvorexant) and tested in 2h sessions.

### Low-effort responding for sucrose reward

A subset of rats (n=12) were trained in 2h sessions to nosepoke for sucrose (45 mg sucrose pellets, Test Diet, Richmond, IN, USA) on an FR1 schedule following locomotor testing. Nosepokes resulted in pellet delivery and were followed by a 20 s time out signaled by termination of the house light. Rats were trained for a minimum of 5 sessions and until the number of pellets obtained differed by less than 25% across 3 consecutive sessions. Suvorexant (0 or 30 mg/kg) was given in a within-subjects, counterbalanced design 30min prior to a subsequent sucrose FR1 test session, with a minimum of 3d between tests.

### Data Analysis

Data are expressed as mean values ± 1 standard error of the mean. Statistics were performed using GraphPad Prism for Mac (Version 7, GraphPad Software Inc., La Jolla, CA) with an α level of 0.05. Qo and α were expressed as percent change from baseline and assessed using separate repeated-measures ANOVAs with Holm-Sidak’s *post hoc* tests. Two-tailed Pearson’s correlations were used to investigate the relationship between baseline demand and suvorexant efficacy. A Kaplan-Meier estimator was used to compare α stabilization rates between suvorexant doses. Separate paired samples t-tests were used to assess changes in suvorexant efficacy, punished responding, and sucrose consumption. Extinction, reinstatement, and locomotor activity data were assessed using mixed-design ANOVAs with Holm-Sidak’s *post hoc* tests.

## Results

### Experiment 1: Efficacy of systemic suvorexant administration on fentanyl demand in rats with limited drug access

The experimental timeline is outlined in Figure 1. Following fentanyl self-administration training, rats (n=19) were trained on a within-sessions BE task and the effects of suvorexant on fentanyl demand were assessed. Figure 2A shows demand curves from a single subject treated with suvorexant (0, 10 and 30 mg/kg) 30min prior to BE testing. Suvorexant pretreatment dose-dependently increased demand elasticity (α; Figure 2B; rm-ANOVA, F_3,54_=5.551, p=0.0021). *Post hoc* analysis revealed that suvorexant at 30mg/kg increased demand elasticity relative to vehicle (Holm-Sidak’s *post hoc* tests, p=0.0026). Vehicle or 10mg/kg suvorexant had no significant effect on demand elasticity (Holm-Sidak’s *post hoc* tests, p=0.1702 and p=0.9422, respectively). Suvorexant had no significant effect on low effort fentanyl consumption (Qo; Figure 2C; rm-ANOVA, F_3,54_=2.226, p=0.0956).

**Figure 1.**
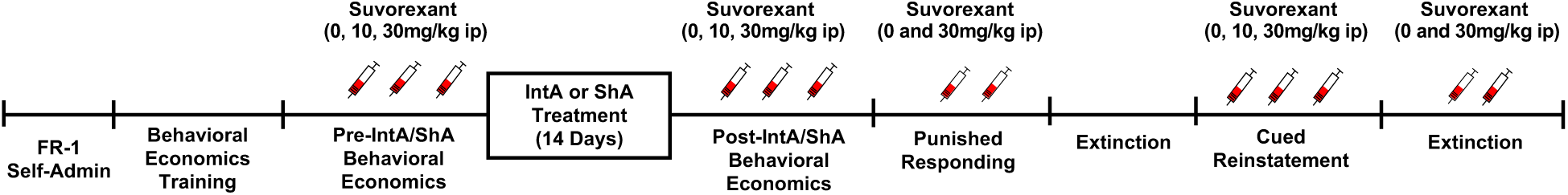
Experimental Timeline. Diagram illustrating the timepoints for behavioral testing

**Figure 2.**
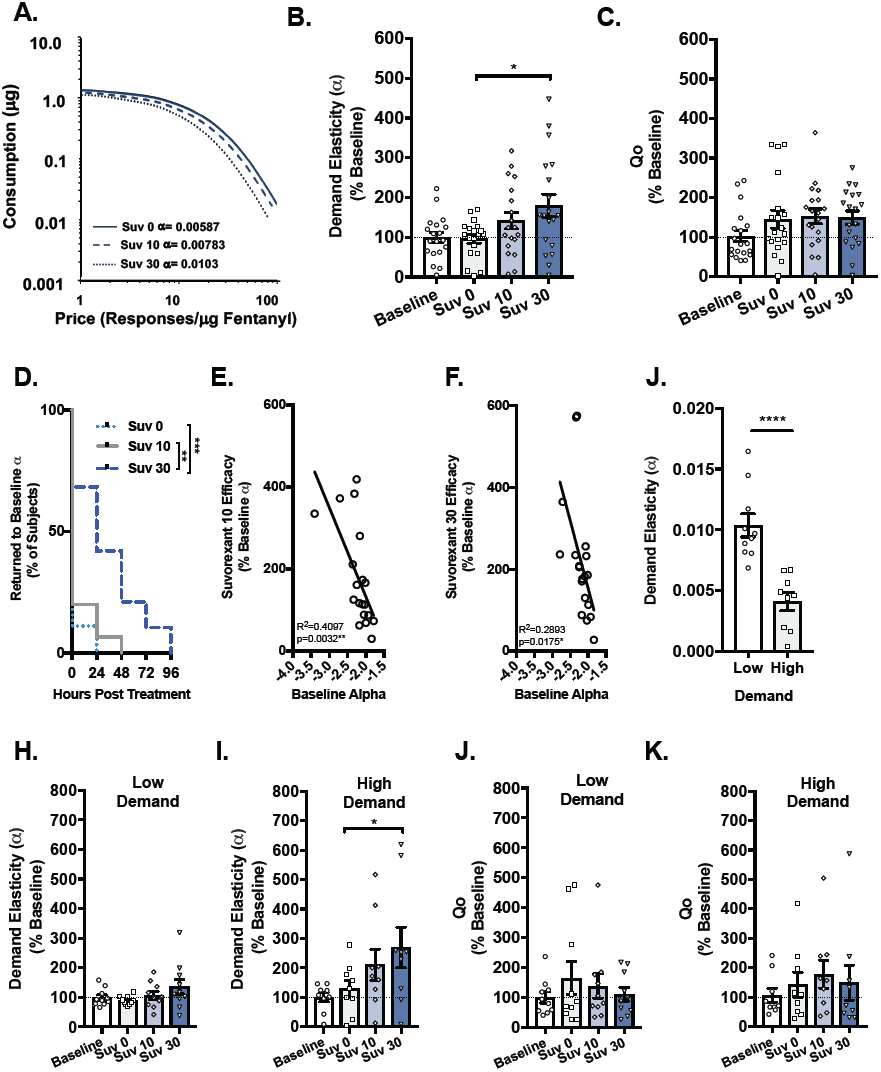
Efficacy of systemic suvorexant administration on fentanyl demand in rats with limited fentanyl access. Rats (n=19) were treated with the OxR1/OxR2 antagonist suvorexant (Suv; 0, 10, or 30mg/kg, ip) prior to BE testing. (A) Shown are demand curves from a single subject that received vehicle (solid) or 10 and 30mg/kg suvorexant (dashed lines). Note the increased demand elasticity (α; represents reduced motivation) after suvorexant. (B) Pretreatment with 30mg/kg suvorexant increased demand elasticity (α) relative to vehicle (rm- ANOVA with Holm-Sidak’s *post hoc* tests, p=0.0026). (C) In contrast, suvorexant had no effect on preferred fentanyl consumption (Qo; rm-ANOVA, p=0.9422). (D) 30mg/kg suvorexant increased demand elasticity (α) for a greater duration compared to vehicle or 10mg/kg suvorexant (Kaplan-Meier estimator, p<0.0001, p=0.0029, respectively). (E-F) Baseline α was negatively correlated with suvorexant (10 and 30mg/kg) efficacy such that suvorexant was most effective in increasing demand elasticity in highly motivated subjects (small α ; Pearson’s r, p=0.0032, p=0.0175, respectively). (G) Rats were separated into low (n=10) and high demand groups (n=9) based on a median split of α values (independent samples t-test, p<0.001). (H) Suvorexant had no effect on demand elasticity in low demand rats (rm-ANOVA, p=0.1294. (I) In high demand rats, suvorexant increased demand elasticity (α) and was most effective at 30mg/kg (rm-ANOVA with Holm-Sidak’s *post hoc* tests, p=0.0352). (J-K) In both low and high demand subjects, suvorexant had no effect on low effort/preferred fentanyl consumption (rm-ANOVA, p=).*p<0.05, **p<0.01, ****p<0.0001

When administered systemically, the OxR1 antagonist SB was previously found to attenuate addiction-like behaviors beyond its bioavailability [15,31]. Here we examined whether suvorexant had similarly persistent effects on fentanyl demand. We observed significantly increased demand elasticity (reduced motivation) at 24h (paired samples t-test; t_17_=2.205, p=0.0415), but not at 48h (t_16_=1.731, p=0.103) or 72h (t_16_=1.276, p=0.223), following 30mg/kg suvorexant treatment (data not shown). There were no persistent effects of suvorexant 10mg/kg (p’s>0.05 on all days). Indeed, survival analyses indicated that α values took significantly longer to return to baseline following 30mg/kg suvorexant compared to vehicle or 10mg/kg suvorexant (Figure 2D; Kaplan-Meier estimator, ϰ^2^_1_=14.72, p<0.0001 and ϰ^2^_1_=8.868, p=0.0029, respectively). The length of time that demand elasticity was reduced did not differ between vehicle and 10mg/kg suvorexant (ϰ^2^_1_=0.959, p<0.959).

We previously demonstrated that baseline demand elasticity (α) for fentanyl predicted the extent to which the OxR1 antagonist SB reduced motivation for fentanyl, such that SB was most effective in rats with inelastic demand (lower α values;[16]). Consistent with these results, baseline α was negatively correlated with the efficacy with which suvorexant increased demand elasticity (Figure 2E-F; 10mg/kg, Pearson’s r, R^2^=0.4097, p=0.0032; 30mg/kg; R^2^=0.2893, p=0.0175). In a subsequent analysis, rats were separated into low- (n=10) and high- (n=9) demand groups based on a median split of baseline α (Figure 3A; independent samples t-test, t_17_=5.088, p<0.001). Qo did not differ between low and high demand groups (data not shown; independent samples t-test, t_17_=0.0215, p=0.9830). Pretreatment with suvorexant had no effect on demand elasticity (α) in low demand rats at either dose (Figure 3C; F_3,27_=2.058, p=0.1294). In high demand rats, suvorexant significantly increased fentanyl demand elasticity (α; Figure 3D; rm-ANOVA, F_3,24_=4.529, p=0.0118) and was most effective at 30mg/kg (Holm-Sidak’s *post hoc* tests, p=0.0352). Suvorexant had no effect on Qo values in low or high demand rats (Figure 2G-H; F_3,27_=0.625, p=0.605 and F_3,24_=0.5064, p=0.6816; respectively).

**Figure 3.**
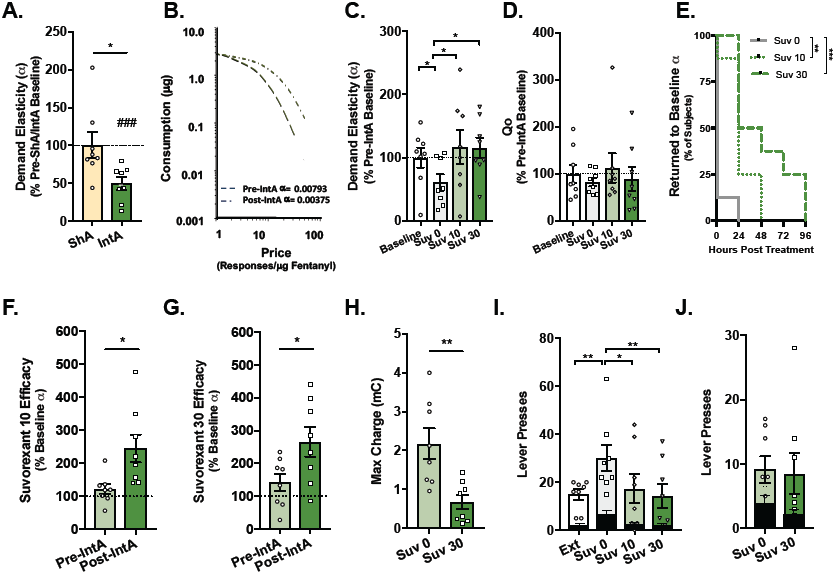
Efficacy of systemic suvorexant on fentanyl demand, punished responding, and cue-induced reinstatement of fentanyl seeking in IntA (addiction model) rats. Rats were assigned to ShA (n=8) or IntA (n=8) groups following initial demand testing. **(A)** IntA to fentanyl was associated with a decrease in demand elasticity relative to baseline α (### denotes significant difference from pre-ShA/IntA baseline, student’s t-test, p=0.0005) and compared to rats given ShA to fentanyl (independent samples t-test, p=0.0174). **(B)** Demand curves from a representative subject before (solid) and after (dashed) IntA to fentanyl. Note reduced demand elasticity (increased motivation) after IntA. **(C)** Prior to BE testing, the suvorexant (Suv; 0, 10 or 30mg/kg, ip) was given to IntA rats. Pretreatment with 10 or 30mg/kg suvorexant increased demand elasticity (α) relative to vehicle (rm-ANOVA with Holm- Sidak’s *post hoc* tests, (p=0.0186 and p=0.0187, respectively). **(D)** Suvorexant had no effect on low effort/preferred fentanyl consumption (Qo; rm-ANOVA, p=0.7931). **(E)** Following IntA to fentanyl, pretreatment with 10 or 30mg/kg suvorexant increased demand elasticity (α) for a longer duration relative to vehicle (Kaplan-Meier estimator, p=0.0037, p=0.0005, respectively). **(F-G)** 10 or 30mg/kg suvorexant were also more effective in increasing demand elasticity (α) after IntA to fentanyl vs. pre-IntA (paired samples t-test, p=0.0186,p=0.0187, respectively). **(H)** When infusions of fentanyl were paired with foot shock of escalating intensity, 30 mg/kg suvorexant decreased the maximal charge accepted by rats to maintain their preferred fentanyl consumption (paired samples t-test, p=0.0048). After testing the effects of suvorexant on fentanyl demand and punished responding, lever press responding was extinguished and the effects of suvorexant on cue-reinstatement were assessed. **(I)** The presentation of fentanyl associated cues increased active lever responding and 10 or 30mg/kg suvorexant attenuated this effect. Suvorexant did not alter responding on the inactive lever (black bars; mixed-design ANOVA with Holm-Sidak’s *post*) **(J)** When administered during extinction, suvorexant did not affect active or inactive (black bars) lever pressing (mixed-design ANOVA). *p<0.05, **p<0.01, ***p<0.001

### Experiment 2: Suvorexant decreases demand, punished responding, and cue-induced reinstatement for fentanyl seeking following IntA to fentanyl

After assessing the therapeutic potential of suvorexant in fentanyl self-administering rats with limited drug access, we sought to investigate the efficacy of suvorexant in an animal model of opioid addiction. Intermittent access (IntA) self-administration induces strong addiction-like behaviors that are orexin-dependent*. Here, we investigated the efficacy of suvorexant in attenuating such addiction-like behaviors following IntA to fentanyl.

Rats were pseudorandomly assigned to short access (ShA) or IntA (n=8 each) following initial BE testing. ShA was included as a comparison to confirm that IntA to fentanyl induced an enhanced addiction-like state. Indeed, IntA to fentanyl was associated with a significant decrease in demand elasticity (α) relative to pre-IntA α (dashed line; Figure 3A; student’s t-test, t_7_=6.0732, p=0.0005), and compared to rats given ShA to fentanyl (independent samples t-test, t_14_=2.697, p=0.0174). Figure 3B shows representative demand curves from an IntA subject illustrating the decrease in demand elasticity following IntA to fentanyl. In IntA rats, pretreatment with suvorexant was effective in increasing demand elasticity (α, decreasing motivation; Figure 3C; rm-ANOVA, F_3,21_=4.004, p=0.0212). *Post hoc* analysis showed that in IntA animals 10mg/kg and 30mg/kg suvorexant increased demand elasticity relative to vehicle (p=0.0186 and p=0.0187, respectively). Suvorexant had no effect on low effort/preferred fentanyl consumption (Qo; Figure 3D; rm-ANOVA, F_3,21_=0.345, p=0.7931). The effect of suvorexant on demand elasticity was persistent; α values were significantly increased at 24h and 48h following both suvorexant 10mg/kg and 30mg/kg (data not shown; paired samples t-test; 10mg/kg, at 24h: t_6_=2.582, p=0.0416; 30mg/kg at 48h: t_4_=3.288, p=0.0303. Indeed, survival analyses revealed that α values took significantly longer to return to baseline following pretreatment with 10 or 30mg/kg suvorexant compared to vehicle (Figure 3E; Kaplan-Meier estimator, ϰ^2^_1_=8.422, p=0.0037 and ϰ^2^_1_=12.47, p=0.0005, respectively). Next, using a within-subjects analysis, we compared the extent to which suvorexant increased demand elasticity (decreased motivation, measure of suvorexant efficacy) before and after IntA self-administration. Both 10mg/kg and 30mg/kg suvorexant were more effective at increasing demand elasticity following IntA (Figures 3F-G; paired samples t-test, t_7_=2.803, p=0.0264; t_7_=2.799, p=0.0266, respectively).

Next we examined the effect of suvorexant on responding for fentanyl when infusions of drug were paired with footshocks of increasing intensity (to measure compulsive responding). To limit the number of footshock sessions in these rats, we tested suvorexant at the 0 and 30mg/kg doses only (2 sessions). Compared to vehicle treatment, suvorexant (30mg/kg) reduced the maximum charge accepted by rats to defend their preferred fentanyl consumption (Figure 3H; paired samples t-test, t_7_=4.063, p=0.0048).

We next assessed the effect of suvorexant on cue-induced reinstatement of extinguished fentanyl seeking in IntA rats. Lever press responding decreased over the first 7 extinction sessions (data not shown; main effect of session F_6,48_ =6.023, p<0.0001). Rats responded more on the active vs inactive lever (main effect of lever type F_1,8_ = 31.51, p=0.0005). However, active vs inactive lever presses were similar by Day 7 of extinction (session x lever type interaction F_6,48_= 4.609, p=0.0009). After reaching extinction criterion, rats were treated with suvorexant (0, 10 or 30 mg/kg) 30min prior to testing for cued reinstatement. We observed significant reinstatement of responding in vehicle-treated rats (Fig. 3I; mixed-design ANOVA; main effect of treatment, F_3,42_=4.189, p=0.0111 and main effect of lever type, F_1,14_=31.33, p<0.001; Holm-Sidak’s *post hoc* tests p=0.0053). This was significantly attenuated by pretreatment with suvorexant at 10 and 30mg/kg (Holm-Sidak’s *post hoc* tests p=0.01 and p=0.0053, respectively). Suvorexant had no effect on inactive lever responding (Fig. 3I, black bars; Holm-Sidak’s *post hoc* tests, p>0.05 all comparisons). To demonstrate that suppression of responding was specific to reinstatement responding, we tested the effect of the highest dose of suvorexant (30mg/kg) on lever responding after extinction was achieved (Figure 3J). Rats responded more on the active vs inactive lever (main effect of lever type F_1,7_ = 7.727, p=0.0273); however was no effect of treatment on active or inactive lever press responding (no main effect of treatment, F_1,7_ = 3.345, p=0.110; no treatment x session interaction F_1,7_ = 0.324, p=0.586).

### Experiment 3: Effect of suvorexant on general locomotion and natural reward seeking

Finally, we sought to determine the effects of the highest dose of suvorexant (30mg/kg) on general locomotor activity and sucrose self-administration, to reveal any non-specific effects on motor activity or natural reward consumption in fentanyl experienced rats. Fentanyl experienced rats received suvorexant (n=10) or vehicle (n=9) 30min prior to locomotor testing. Suvorexant had no effect on the total distance traveled in the novel locomotor chamber over the 2h test session (Figure 4A; mixed-design ANOVA; no effect of treatment F_1,17_=0.6645, p=0.4263 or treatment x time interaction: F_1,17_=0.6645, p=0.4263). Following locomotor testing, a subset of fentanyl experienced rats (n=12) were trained to self-administer sucrose pellets on an FR1 schedule. Suvorexant pretreatment also had no effect on the number of sucrose pellets earned during a 1h self-administration session (Figure 4B; paired-samples t-test, t_11_=1.748,p=0.1083). Together, the data indicate that general locomotor activity and lever pressing for a non-drug reinforcer were not impaired by suvorexant treatment, and that suvorexant effects on addiction-like behavior were not due to a motor impairment.

**Figure 4:**
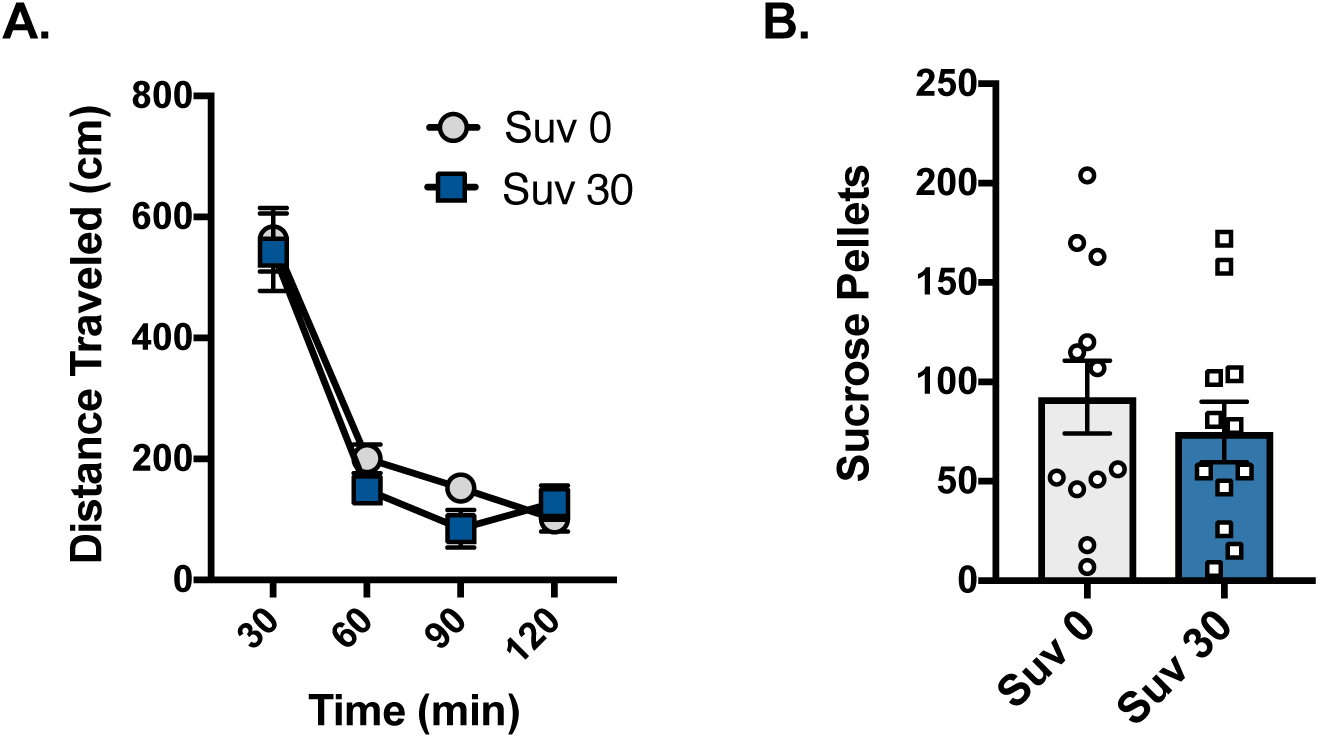
Effect of suvorexant on general locomotion and sucrose consumption. **(A)** In fentanyl experienced rats (n=19), suvorexant (Suv; 30 mg/kg, i.p.) had no effect on the total distance traveled in a novel locomotor environment over the 2h session when assessed in 30min bins (mixed-design ANOVA, p=0.4263). **(B)** 30 mg/kg suvorexant also had no effect on sucrose consumption in rats (n=12) trained to self-administer fentanyl by nose-poking on an FR-1 schedule (paired samples t-test, p=0.1083).

## Discussion

The present study investigated the therapeutic potential of the FDA-approved dual OxR1/OxR2 antagonist suvorexant for treating addiction-like behaviors in fentanyl self-administering rats. We found that systemic suvorexant increased demand elasticity (decreased motivation) for fentanyl without affecting preferred fentanyl consumption at null cost (Qo). Suvorexant was most effective at increasing demand elasticity in rats with the highest baseline motivation for fentanyl; at the highest dose tested, suvorexant increased demand elasticity for up to 24h post-treatment.

IntA to fentanyl increased demand for fentanyl across all rats, as we previously reported [12]. Suvorexant more effectively increased demand elasticity, and increased demand elasticity for a longer duration, after IntA compared to ShA treatment. Suvorexant also reduced punished responding for fentanyl and cue-induced reinstatement of fentanyl seeking. Suvorexant did not alter extinction responding, general locomotor activity, or sucrose self-administration, indicating that the actions of suvorexant to decrease addiction-like behaviors were not due to sedation, attenuated motor ability or nonspecific decreases in reward consumption. These results indicate that suvorexant may be an effective treatment for OUD and may best benefit those with the most serve cases of opioid addiction.

### A role for OxR1 and OxR2 signaling in opioid addiction

In rodent models, OxR1 antagonists reliably decrease the reinforcing and relapsing properties across multiple drugs of abuse [34-39], particularly in drug seeking behaviors that require high effort or during enhanced addiction-like states[16,17,29,40-42]. Recently, we demonstrated that the selective OxR1 antagonist SB increased fentanyl demand elasticity (decreased motivation) in rats with limited drug access, an effect that was greatest in highly motivated rats[16]. We also found that highly motivated rats after IntA to fentanyl are more sensitive to the anti-addiction properties of SB compared to rats given ShA [12] a finding that is consistent with evidence that SB preferentially reduces motivation for other drugs of abuse in high-motivation individuals[14,17,42-44]. Here, we show that suvorexant has anti-addiction properties similar to the selective OxR1 antagonist, SB. In rats with limited drug access, we found that suvorexant preferentially increased fentanyl demand elasticity in rats with the greatest motivation for fentanyl. Following IntA, we found that suvorexant decreased motivation for fentanyl at a lower dose (10mg/kg) than was effective before IntA. We also showed that suvorexant is more effective in decreasing motivation for fentanyl after IntA and that suvorexant attenuates other addiction indices including compulsive responding for fentanyl and cued reinstatement of fentanyl seeking.

Growing evidence supports a role for OxR2 signaling in opioid reward and dependence. For example, intra-VTA administration of the selective OxR2 antagonist TCS-Ox229 blocked the acquisition and expression of morphine CPP [45] as well as stress- and prime-induced reinstatement of extinguished morphine CPP [46]. Moreover, local injections of TCS-Ox229 into paraventricular thalamus blocked the expression of naloxone-induced conditioned place aversion [47]. OxR2 signaling may also regulate drug taking in situations of enhanced motivation. Systemic administration of the OxR2 antagonist NBI-80713 reduced heroin self-administration in highly motivated rats after long (LgA), but not after ShA, to heroin [20]. LgA to heroin was also associated with an increase in OxR2 mRNA in critical brain stress structures [20], indicating that OxR2 signaling becomes particularly important during enhanced ‘addiction-like’ states like that induced in rats given IntA to fentanyl. Together, these findings indicate that dual OxR1/OxR2 antagonists, including suvorexant, might have more profound anti-addiction properties compared to selective OxR1 antagonists, particularly in highly addicted individuals. To this end, it will be important that future studies directly compare the efficacy of single- vs. dual-OxR antagonists, ideally using a model (such as the behavioral economics procedure used here) that allows for repeated pharmacological testing in the same animal.

### Persistent effects of suvorexant on fentanyl demand

Our laboratory and others have shown that systemic administration of the OxR1 antagonist SB attenuates drug seeking beyond the bioavailability of the compound [15,31]. Consistent with this, here we report that in rats with limited drug access, 30 mg/kg suvorexant increased fentanyl demand elasticity for ∼24h. Moreover, suvorexant increased demand elasticity for a longer duration following IntA, and this effect occurred at lower suvorexant doses. In rats, suvorexant has a relatively short half-life of ∼0.6h [48], indicating that the persistent effects seen here stem from neuroadaptations in downstream signaling pathways involved in motivation [31]. These results have important implications concerning the optimal dosing regimen for suvorexant in OUD treatment. For example, every other day dosing of suvorexant may be sufficient to suppress craving but also reduce the risk of sedation. Clearly, future studies are required to test this possibility.

### Suvorexant was not sedating at doses that suppressed drug seeking

When administered at bedtime, suvorexant reduces latency to sleep onset and promotes longer sleep time[48]. Thus, sedation is a potential concern with repurposing suvorexant for OUD. Notably, we found no evidence that suvorexant produced soporific effects at doses that suppressed fentanyl seeking when administered during the active period. Suvorexant had no effect on locomotor activity in a novel open field or on responding for sucrose on an FR1 schedule, nor did it affect responding on an extinction test. Our results are consistent with previously published studies demonstrating that, at the doses tested here, suvorexant does not alter cocaine-induced hyperlocomotion or general responding for sucrose reward on a 5-choice serial reaction time task [24,49]. Although there is some evidence that 30mg/kg suvorexant modestly decreases homecage locomotor activity in rats [48], the homecage environment lacks the sensory stimulation that was present during the novel locomotor and operant tasks used here, and thus suvorexant may be less likely to promote somnolence in stimulating environments. An additional consideration is the fact that chronic opioid administration augments the orexin system; orexin protein levels are increased in rodents chronically exposed to fentanyl or morphine [11,12] and morphine reverses cataplexy in mice made narcoleptic through orexin depletion [11]. Consequently, the doses of suvorexant that effectively reduce drug craving may differ from those that induce sedation.

### Leveraging the sleep-promoting effects of suvorexant to reduce relapse risk

Sleep impairments are common in OUD, including insomnia, disrupted sleep architecture, as well as sleep-related breathing disorders [21,50]. Sleep disturbances are also common in patients receiving OST, such as methadone and buprenorphine [51]. Poor sleep quality might negatively impact outcomes and adherence to addiction treatments, and impaired sleep promotes poor executive function and negative affect, which both increase relapse risk [52,53]. Thus, in addition to reducing the activity of hyperexctiable brain circuits that promote motivated reward seeking, it is possible that suvorexant may have additional therapeutic properties in OUD patients by improving sleep quality and leading to better emotional and executive functioning[21]. However, the sleep promoting effects of suvorexant are achieved when administered immediately prior to sleep onset, which is distinct from the dosing regimen used here where suvorexant was administered during the active period (when drug opportunity/use is highest in OUD patients). It is not clear whether the anti-drug seeking properties of suvorexant would persist into the active period if administered prior to sleep, however our finding that suvorexant reduced demand for fentanyl beyond its bioavailability (discussed above) indicates that this might be the case. This is an important avenue for future studies on this topic.

### Conclusions

We show that the dual OxR1/OxR2 receptor antagonist suvorexant effectively decreases fentanyl demand and other addiction-related behaviors with no observable off-target effects. Notably, suvorexant was most effective in attenuating addiction behaviors in the most highly motivated rats. Together, our findings provide initial support for the use of suvorexant in the treatment of OUD, particularly in those individuals with the most severe cases of opioid addiction.

